# Characterization of A Bronchoscopically Induced Transgenic Lung Cancer Pig Model for Human Translatability

**DOI:** 10.1101/2024.11.04.621940

**Authors:** Kirtan Joshi, Kanve N. Suvilesh, Nagabhishek Sirpu Natesh, Yariswamy Manjunath, Jared Coberly, Sarah Schlink, Jeffrey R. Kunin, Randall S. Prather, Kristin Whitworth, Benjamin Nelson, Jeffrey N. Bryan, Timothy Hoffman, Mojgan Golzy, Murugesan Raju, Emma Teixeiro, Bhanu P. Telugu, Jussuf T. Kaifi, Satyanarayana Rachagani

## Abstract

**Background:** There remains a need for animal models with human translatability in lung cancer (LC) research. Findings in pigs have high impact on humans due to similar anatomy and physiology. We present the characterization of a bronchoscopically-induced LC model in Oncopigs carrying inducible KRAS^G12D^ and TP53^R167H^ mutations.

**Methods:** Twelve Oncopigs underwent 29 injections via flexible bronchoscopy. Eighteen Adenovirus-Cre recombinase gene (AdCre) inductions were performed endobronchially (n=6) and transbronchially with a needle (n=12). Eleven control injections were performed without AdCre. Oncopigs underwent serial contrast-enhanced chest CT with clinical follow-up for 29 weeks. Following autopsy, lung and organ tissues underwent histopathology, immunohistochemistry, and RNA-sequencing with comparative analysis with The Cancer Genome Atlas (TCGA) human LC data.

**Results:** All 18 sites of AdCre injections had lung consolidations on CT imaging. Transbronchial injections led to histopathologic invasive cancer and/or carcinoma in situ (CIS) in 11/12 (91.7%), and invasive cancer (excluding CIS) in 8/12 (66.6%). Endobronchial inductions led to invasive cancer in 3/6 (50%). A soft tissue metastasis was observed in one Oncopig. Immunohistochemistry confirmed expression of Pan-CK+/epithelial cancer cells, with macrophages and T cells infiltration in the tumor microenvironment. Transcriptome comparison showed 54.3% overlap with human LC (TCGA), in contrast to 29.88% overlap of KRAS-mutant mouse LC with human LC.

**Conclusions:** The transgenic and immunocompetent Oncopig model has a high rate of LC following bronchoscopic transbronchial induction. Overlap of the Oncopig LC transcriptome with human LC transcriptome was noted. This pig model is expected to have high clinical translatability to the human LC patient.

## BACKGROUND

Animal models, with a majority being mice ^1^, are crucial in advancing diagnostic and therapeutic discoveries to human lung cancer (LC) patient care. However, most of the treatments developed in mice have failed in clinical trials, resulting in losses in capital, time, and patient lives ^2^. While mice do not adequately mimic human patients ^3–5^, pigs share many similarities with humans including a relatively longer life span, body size, anatomy, physiology, diet, metabolome, immune system, and genetics ^3, 4, 6, 7^. Research in pigs resulted in major therapeutic advancements of human diseases, such as cystic fibrosis ^8^. Testing in pig models allows short- and long-term trialing of human-grade novel medical/surgical devices, imaging/radiation/theranostics technologies, and drugs^9^.

Oncopigs are a transgenic pig line carrying the common tumor suppressor gene TP53^R167H^ and oncogene KRAS^G12D^ mutations that are inducible in any organ of choice utilizing a *Cre/Lox* system ^10–12^. Both mutations are orthologous to the mutations observed in human cancers ^11–14^. While TP53 is the most commonly mutated tumor suppressor gene in human LC ^15^, KRAS is the most frequently mutated oncogene in non-small cell lung cancer (NSCLC) adenocarcinoma ^16^. The immunocompetent Oncopig model carrying those two predominant and deterministic driver mutations of cancers offers outstanding opportunities to study LC through lung-specific, targeted induction via Adenoviral-mediated Cre recombinase (AdCre) delivery ^17, 18^. In previous studies, Oncopig liver and pancreatic cancers were induced via organ-specific induction with AdCre, also demonstrating striking similarities with the matched human cancers ^19, 20^.

A recent pilot study on four Oncopigs led to histopathologically confirmed LC induction at a maximum rate of 33.3% via percutaneous injection with a surveillance period of 21 days post-induction ^21^. To build on this pilot study and increase LC induction rates ^21^, we chose a larger cohort of Oncopigs and performed AdCre injections via flexible bronchoscopy with an extended surveillance period of up to 29 weeks. High-resolution, contrast-enhanced CT chest imaging showed consolidations at all injection sites, but none in the controls. Upon autopsy, invasive LC and distant metastatic disease were confirmed histopathologically as PanCK+ cells, with transbronchial injections leading to the highest LC and carcinoma in situ (CIS) induction rates of >90%. LCs showed substantial immune cell infiltration in the tumor microenvironment.

Comparison with orthologous human LC gene expression data from The Cancer Genome Atlas (TCGA) demonstrated much higher concordance of transgenic pig LC than mouse LC. It is anticipated that an LC Oncopig model carrying human patient-relevant mutations and substantial transcriptome overlap will leverage translational LC research findings for human LC patients and, ultimately, improve their outcomes.

## MATERIALS AND METHODS

### Pigs and bronchoscopic tumor induction in the lung

The study was approved by the Institutional Animal Care and Use Committee at the University of Missouri (protocol # 18120). Animal housing facilities are accredited by AAALAC and in compliance with and regulated by USDA (43R0048) and Office of Laboratory Animal Welfare (D16-00249). Generation of the Oncopig line is described in School et al ^10^. 9-14-week-old Minnesota mini Oncopigs (LSL-KRAS^G12D^-IRES-TP53^R167H^) were acquired from the National Swine Research and Resource Center (NSRRC) (Columbia, MO) and transported to the Animal Sciences Research Center (ASRC) at the University of Missouri, Columbia. The animals were allowed to acclimate to the facility for 3 days before any experimental intervention.

### Endo- and transbronchial injections of Adenovirus-Cre (AdCre)

Ad5CMVCre-eGFP (high titer) virus was obtained from the University of Iowa Viral Vector Core. A concentration of 1×10^11^ plaque-forming units (PFU)/mL was used for group 1 and 2. A volume of 150 µL of AdCre (1×10^11^ PFU/mL) ± adjuvants porcine IL-8 (5 ng/mL) (NBP2-35234; Novus Biological) and ± adjuvants Polybrene (1:100) were injected. In group 3, we chose to give a 3x higher dose (3×10^11^ PFU/mL) of AdCre as had been suggested for development of a pancreatic cancer transgenic Oncopig model^22^. The rationale to use cytokine IL-8 with AdCre is that it mobilizes the adenovirus receptor to the luminal membrane of epithelial cells, thereby enhancing viral entry ^22–24^. Polybrene (hexadimethrine bromide) was used to increase binding between pseudoviral capsid and the cellular membrane to enhance transduction efficiency ^25^.

Anesthesia and procedures were performed or supervised by veterinary physicians with expertise in clinical veterinary care, a general thoracic surgeon with expertise trained in airway management and interventional bronchoscopy, and ancillary veterinary staff trained in general pig care. Pigs were anesthetized using ketamine and xylazine. During interventions, the pigs were placed in laterally recumbent position and maintained under general anesthesia using inhalation isoflurane with either a nose cone or a laryngeal mask. Pigs were anesthetized for a relatively short time during the bronchoscopic inoculation and during the CT scans. The average time under anesthesia was approximately 15 minutes or shorter.

Bronchoscopic injections were performed via laryngeal mask using an Ambu aScope^TM^ 4 Bronchoscope (regular, 5.0/2.2) that was inserted transorally. An Interject^TM^ Injection Therapy needle catheter (25-gauge, 6 mm needle) was used to inject transbronchially. Injections via flexible bronchoscopy were done with two techniques: (1) Endobronchial or (2) Transbronchial needle injection into different lung lobes. As controls, adjuvants (polybrene ± IL-8) or vehicle [phosphate-buffered saline (PBS)] alone without AdCre were injected into the contralateral lung lobes in the same pig. The bronchoscope was passed through a laryngeal mask and navigated to the target lobe (**Table 1**). The inoculation mixtures were either injected endobronchially via the bronchoscopy channel or transbronchially using the needle (**Figure 1**). For transbronchial injections, the injection needle catheter dead space was flushed with 200 µL PBS to ensure that all AdCre was inoculated.

**Figure 1:**
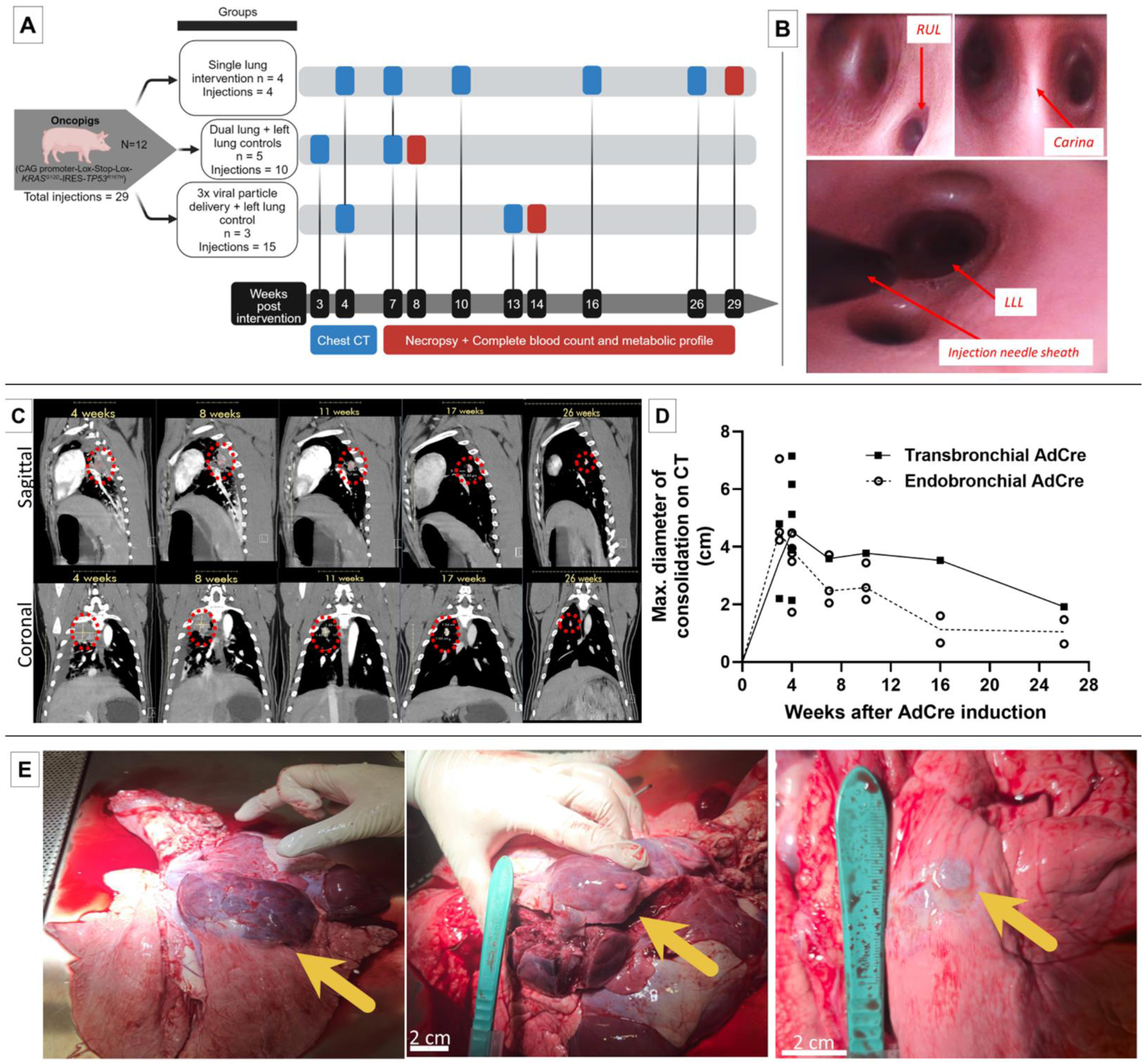
**A:** Study design: Bronchoscopic inductions (total n=29) were performed in twelve Oncopigs separated into three groups. Clinical surveillance and follow-up imaging were performed with contrast-enhanced chest CT at the given time points. Blood analyses and necropsies were performed a three different time points post-induction. **B**: **Flexible bronchoscopy images from Oncopig airways.** Left upper panel: Right upper lobe (RUL) bronchus that is located proximally to the main carina. Right upper panel: Close-up view of the main carina with bifurcation into right and left main stem bronchus. Lower panel: Bronchoscopic injection needle sheath visible in the left lower lung lobe (LLL). All interventions in pigs were performed with human-grade bronchoscopy equipment. **C: High-resolution, contrast-enhanced chest CT scans following bronchoscopic AdCre induction in Oncopigs.** Sagittal and coronal images showing consolidation in the lung at the AdCre injection sites at various timepoints. Regression of the consolidations were observed over subsequent imaging, with persistence of solid lung nodules. Contralateral lung lobe injections with adjuvant agents (IL-8, polybrene) without AdCre did not show any consolidations on CT imaging. **D: Curves showing radiographic consolidation on CT imaging** determined by maximum consolidation diameter (cm) measurements according to the AdCre injection techniques (endo-versus transbronchial). (Connecting line: medians). **E**: **Autopsy images from Oncopigs** that had undergone LC induction with bronchoscopic injection of AdCre. Left panel: Right upper lobe lung (RUL) mass; Middle panel: Same dissected tumor; Right panel: Right upper lobe (RUL) nodule. (Arrows indicate tumors).

**Table 1:** Oncopig characteristics, injection/procedure details, and histopathology findings.

| Group | Pig<br>[#] | Sex | Age<br>[Weeks] | Weight<br>[kg] | Injection<br>Number | Injection<br>Method | Injection<br>Agents | Lung Injection<br>Location |  | Histopathology<br>[Invasive cancer or CIS] |
| --- | --- | --- | --- | --- | --- | --- | --- | --- | --- | --- |
|  |  |  |  |  |  |  |  | Side | Lobe |  |
| 1 | 200 | M | 9 | 28 | 1 | Endobronchial | <u>AdCre</u> + IL-8 + PB | Right | Upper | No |
| 1 | 191 | F | 9 | 29 | 2 | Endobronchial | <u>AdCre</u> + IL-8 + PB | Right | Upper | No |
| 1 | 192 | F | 9 | 24 | 3 | Transbronchial | <u>AdCre</u> + IL-8 + PB | Right | Upper | No |
| 1 | 193 | F | 9 | 27 | 4 | Endobronchial | <u>AdCre</u> + IL-8 + PB | Right | Upper | No |
| 2 | 12-2 | F | 9 | 29 | 5 | Transbronchial | <u>AdCre</u> + IL-8 + PB | Right | Upper | Yes |
| 2 |  |  |  |  | 6 | Transbronchial | IL-8 + PB | Left | Lower | No |
| 2 | 12-5 | F | 9 | 32 | 7 | Endobronchial | <u>AdCre</u> + IL-8 + PB | Right | Upper | Yes |
| 2 |  |  |  |  | 8 | Endobronchial | IL-8 + PB | Left | Lower | No |
| 2 | 12-6 | F | 9 | 33 | 9 | Endobronchial | <u>AdCre</u> + IL-8 + PB | Right | Upper | Yes |
| 2 |  |  |  |  | 10 | Endobronchial | PB | Left | Lower | No |
| 2 | 12-8 | M | 9 | 34 | 11 | Endobronchial | <u>AdCre</u> + IL-8 + PB | Left | Lower | Yes (+ soft tissue metastasis) |
| 2 |  |  |  |  | 12 | Endobronchial | IL-8 | Right | Upper | No |
| 2 | 12-9 | M | 9 | 32 | 13 | Transbronchial | <u>AdCre</u> + IL-8 + PB | Right | Upper | Yes |
| 2 |  |  |  |  | 14 | Transbronchial | PB | Left | Lower | No |
| 3 | 54-4 | F | 14 | 33 | 15 | Transbronchial | <u>AdCre</u> + IL-8 | Right | Upper | Yes |
| 3 |  |  |  |  | 16 | Transbronchial | IL-8 | Right | Middle | No |
| 3 |  |  |  |  | 17 | Transbronchial | PBS buffer only | Left | Lower | No |
| 3 |  |  |  |  | 18 | Transbronchial | <u>AdCre</u> + IL-8 | Left | Upper | Yes |
| 3 |  |  |  |  | 19 | Transbronchial | <u>AdCre</u> only | Left | Lower | Yes |
| 3 | 54-8 | F | 14 | 36 | 20 | Transbronchial | <u>AdCre</u> + IL-8 | Right | Upper | Yes |
| 3 |  |  |  |  | 21 | Transbronchial | <u>AdCre</u> + IL-8 | Right | Middle | Yes |
| 3 |  |  |  |  | 22 | Transbronchial | <u>AdCre</u> only | Right | Lower | Yes |
| 3 |  |  |  |  | 23 | Transbronchial | IL-8 | Left | Upper | No |
| 3 |  |  |  |  | 24 | Transbronchial | PBS buffer only | Left | Lower | No |
| 3 | 54-12 | M | 14 | 34 | 25 | Transbronchial | <u>AdCre</u> + IL-8 | Right | Upper | Yes [carcinoma in situ (CIS)] |
| 3 |  |  |  |  | 26 | Transbronchial | <u>AdCre</u> + IL-8 | Right | Middle | Yes (CIS) |
| 3 |  |  |  |  | 27 | Transbronchial | <u>AdCre</u> only | Right | Lower | Yes (CIS) |
| 3 |  |  |  |  | 28 | Transbronchial | IL-8 | Left | Upper | No |
| 3 |  |  |  |  | 29 | Transbronchial | PBS buffer only | Left | Lower | No |

Abbreviations: AdCre: adenovirus Cre recombinase; CIS: carcinoma in situ; F, female; IL-8: interleukin-8; M, male; PB: polybrene; PBS: phosphate-buffered saline

### Computed tomography imaging

Oncopigs were clinically monitored daily for pain and distress including coughing, tachypnea, lethargy, or weight loss was maintained. If excessive coughing was witnessed in the pigs, 3.75mg/kg of Carprofen was administered daily for 5 days to the Oncopigs. For contrast-enhanced computed tomography (CT) (CT scanner: Definition Edge 128 slice; Siemens) 2 mL/kg Omnipaque (350 mg/mL) was injected intravenously with a 15 mL saline flush. The rate of injection was 2 mL/sec with a 40-second delay between initiating the contrast injection and scan. All CT imaging was stored in a DICOM medical image management system (Ambra) and interpreted with a written report by a board-certified radiologist specialized in thoracic imaging.

### Blood draws, autopsy, pathological analysis and sequencing of tissues

At the time of autopsy, a minimum 10 mL of blood was collected via a jugular vein phlebotomy and collected in K2-EDTA and heparin tubes (BD Vacutainer). Complete blood counts (CBCs) and basic metabolic panels (BMPs) were determined. Animals were euthanized using an intravenous Euthasol injection (1 mL/4.5 kg; Virbac).

Following euthanasia, autopsy with gross exam of all major organs including thoracic and abdominal lymph nodes was performed under supervision of a veterinary pathologist. Tissues were fresh-frozen or formalin-fixed for histopathology. Following processing in alcohol and saline, tissues were embedded in paraffin and sectioned at 5 µm thickness. Slides were stained for hematoxylin & eosin and immunohistochemistry was performed with antibodies for Pan-Cytokeratin (Pan-CK) (MNF116, mouse; Agilent), Ionized Calcium-Binding Adaptor molecule 1 (IBA-1) (ab5076, goat; Abcam), and CD3 (ab135372, rabbit; Abcam). Additional immunostaining was performed using E-cadherin (3195S, rabbit; CST), Ki-67 (9449S, mouse; CST), and vimentin (V6630, mouse; Sigma). Chromogen was 3,30-diaminobenzidine tetrachloride (DAB) and the counterstain was hematoxylin. Masson’s Trichrome staining was performed to visualize collagen in tissues. All immunohistochemical staining was validated using appropriate swine control tissues serving as negative controls. Slides were then evaluated and scored blinded by a board-certified pathologist. RNA-sequencing of pig tissues are described in the **Supplementary Methods**.

### Statistical analysis

Statistical difference between more than two groups has been calculated via contingency table analysis with Fisher’s exact test. All standard statistical analysis was done using Prism v8.00 (GraphPad). Bioinformatic analyses of sequencing data and cross species orthologous gene expression analyses are outlined in the **Supplementary Methods**. A *P-value* of <0.05 was considered significant.

## RESULTS

### Bronchoscopic lung cancer induction in Oncopigs

Twelve Oncopigs [n=12; 8 (66.6%) females, and 4 (33.3%) males] were included for LC induction via flexible bronchoscopy (details provided in **Table 1**). Injections were performed in three cohorts in pigs 9-14 weeks of age: Group 1. Single lung intervention (n=4 pigs; autopsied week 29), Group 2. Dual lung + left lung controls (n=5 pigs; autopsied week 8), Group 3. Increased Adenovirus-Cre particle delivery (3x the quantity than in group 1 & 2) + left lung control (n=3 pigs; autopsied week 14) (**Figure 1A**). Slight variations in use of adjuvants polybrene and IL-8, and increased AdCre dose in group 3 were used to adapt to findings in the 1^st^ cohort (e.g., histopathological lack of invasive cancer in group 1, inflammatory reactions observed in group 1&2). In 12 Oncopigs, a total of 29 bronchoscopic injections were performed, out of these 9 (31.0%) endobronchially and 20 (69%) transbronchially with a needle (**Figure 1B**). 18 (62.1%) LC induction attempts with AdCre were performed. 11 (37.9%) control injections were done without AdCre, but with adjuvants polybrene ± IL-8 or phosphate-buffered saline (PBS) only. AdCre was injected endobronchially [n=6/18 (33.3%)] and transbronchially [n=12/18 (66.7%)] into different lung lobes (**Table 1**). No complications, such as endobronchial bleeding or pneumothorax, were encountered with the transbronchial needle injection.

### Clinical surveillance and high-resolution chest CT imaging following induction

Oncopigs were monitored clinically and with high-resolution, contrast-enhanced chest computed tomography (CT) imaging at several time points until autopsy (**Figure 1A**). All pigs remained healthy with expected weight gains with normal CBCs and BMPs at the time of autopsy. All 18 (100%) AdCre injection sites led to radiographic lung consolidations at the injection site on the first contrast-enhanced chest CT at 3 weeks (**Figure 1C**). None of the 11 control injection sites without AdCre showed any consolidations. On CT imaging, consolidations in the lung were peaking in size at the first imaging at 3 weeks [Endobronchial: mean 5.27 cm (standard deviation (SD) ±1.56), median 4.52 cm (range 4.23-7.06 cm); Transbronchial: mean 3.50 cm (SD ±1.84), median 3.50 cm (range 2.20-4.80 cm)], subsequently regressing while there remained persistent nodules radiographically (**Figure 1D**). Of note, one Oncopig developed a pleural effusion that resolved. In some Oncopigs, a rather mild mediastinal lymphadenopathy was observed on CT that was interpreted as reactive. At the latest radiographic time point at 26 weeks, lung consolidations from endobronchial injections [mean 1.05 cm (SD ±0.60), median 1.05 cm (range 0.63-1.47 cm)] were smaller than the transbronchial injection [1.92 cm (n=1)]. No signs of organ metastases were observed on CT imaging.

### LC induction rates are highest with transbronchial injection with higher AdCre dose

At the time of autopsy (group 1: 29 weeks, group 2: 8 weeks, group 3: 14 weeks), all 18 (100%) AdCre injection sites had macroscopic masses in the lungs (**Figure 1E**). There was no macroscopic evidence of metastasis in the locoregional or distant lymph nodes. All other organs inspected (including brain, liver, adrenals, kidneys) were normal, except an incidental finding of soft tissue mass in one Oncopig (12-8). None of the 11 control injection sites revealed macroscopic lung tumors on autopsy.

Tumors, adjacent healthy-appearing lung, soft tissue metastasis and multiple organ tissues were sectioned and stained. Hematoxylin & eosin and immunohistochemistry staining was done to identify invasive, Pan-CK+ LC cells (**Figures 2-4**; **Table 1**). Out of the total 18 endo-& transbronchial AdCre injection sites, 11 (61.1%) revealed invasive cancer cells and 3 (16.7%) carcinomas in situ (CIS), i.e., 14/18 (77.8%) injection sites had CIS or invasive cancer (**Figure 2A**; **Table 1**). Cancers appeared undifferentiated non-small cell lung cancers (NSCLC) without typical histological pattern of an adeno- or squamous cell carcinoma NSCLC, and no small cell carcinoma (SCLC) features were identified histopathologically.

**Figure 2:**
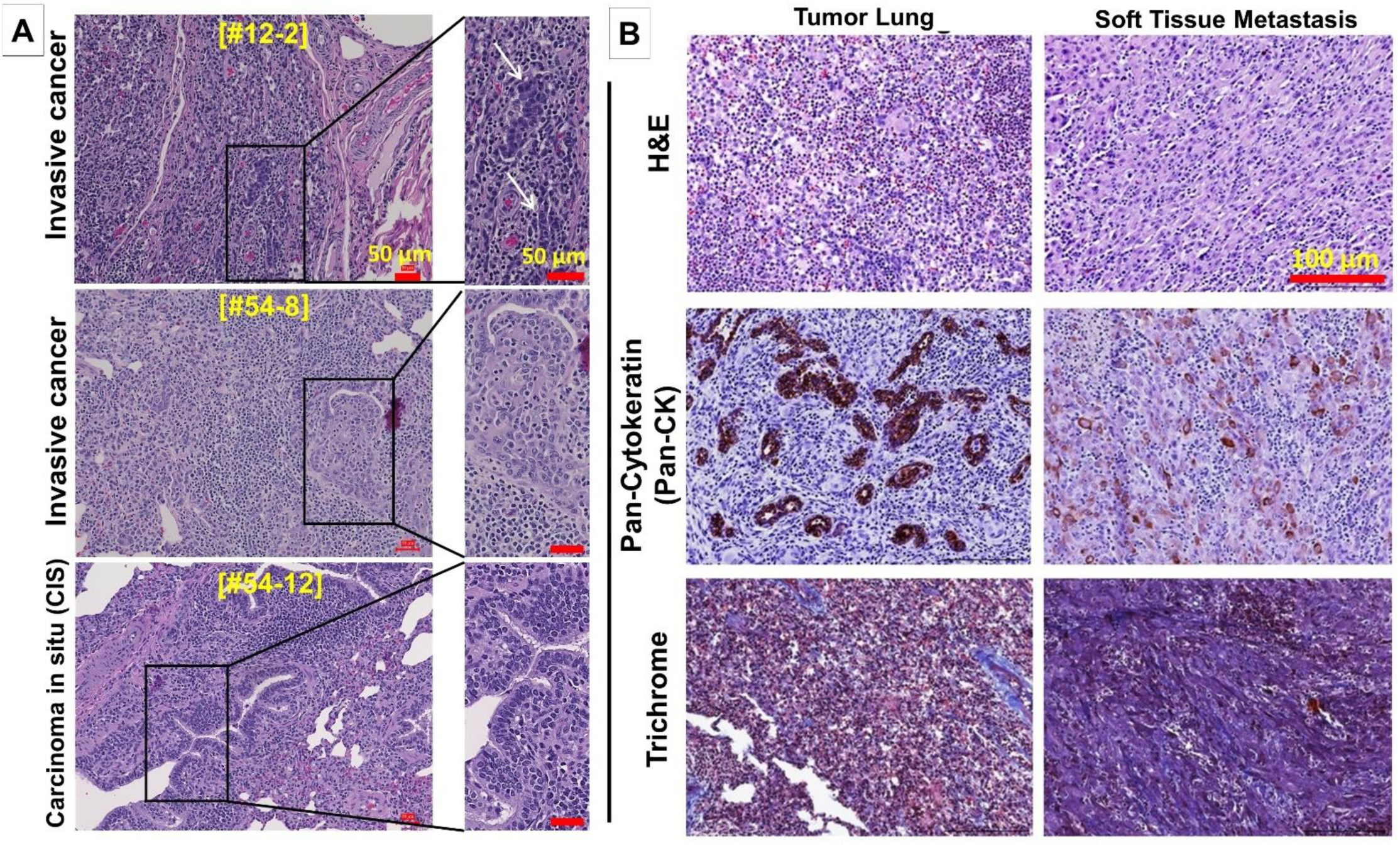
**A:** Hematoxylin and eosin (H&E) staining of three different Oncopig lung tumors showing invasive cancer cells (upper and middle panels) and carcinoma in situ (CIS) (lower panel). (Scale bars, 50 μm). **B: Histopathology and immunostaining** of matched Oncopig LC (left panels) and soft tissue metastasis (right panels) stained with H&E (top row), Pan-Cytokeratin (Pan-CK) for epithelial cells (middle row), and Masson’s Trichrome for visualization of connective tissue/stroma (lower row). (Scale bars, 150 μm).

Transbronchial injections were more efficient in LC induction and led to CIS or invasive cancer in 11/12 (91.7%), and to invasive cancer (excluding CIS) in 8/12 (66.6%). In contrast, endobronchial AdCre injections led to invasive cancer in 3/6 (50%), with no detection of CIS only. Comparing induction of invasive cancer and CIS, transbronchial injections appeared more successful in contrast to endobronchial injections, although not reaching level of significance (P=0.083; Fisher’s exact test). No CIS/cancer was found at any of the 11 control injection sites, and no cancer cells were observed in any of the adjacent healthy lung tissues, draining hilar/mediastinal lymph nodes, or other organs (**Table 1**). One Oncopig had a soft tissue metastasis consisting of invasive, Pan-CK+ cancer cells (**Figure 2B**). Of note, group 3 (that received 3x higher AdCre delivery load in comparison to group 1&2) showed in 6/9 (66.7%) invasive cancer at the AdCre injection sites. In the remaining 3/9 (33.3%) AdCre injection sites CIS was noted (**Figure 2A, lower panels**). This higher induction rate indicated that the AdCre load, combined with transbronchial injection, may contribute to higher LC induction rates. Of note, Masson’s trichrome stain showed moderate presence of connective tissue in the primary LCs and soft tissue metastasis (**Figure 2B**).

### Immune cell infiltration in the Oncopig LC tumor microenvironment

Significant immune cell infiltration, including histiocytes and some granulomata, was found histopathologically on H&E staining in all 18 AdCre injection sites. Immune cell-specific immunohistochemistry identified macrophages (IBA-1+) and T cells (CD3+) in the tumor microenvironment (**Figure 3A/B**). **Figure 3A** shows immunohistochemistry for epithelial cells (Pan-CK), macrophages (IBA-1+) and T cells (CD3) from all three groups with autopsy dates at 8, 14, and 29 weeks. Although these three groups were induced with slight variations, these different autopsy dates suggested initially abundant immune cell infiltration early at 8 weeks following AdCre injection at the tumor sites, decreasing at weeks 14 and 29. Consistent with the regression of the consolidation noted on CT imaging, the overall quantity of immune cell infiltration in the tumor microenvironment decreased with a longer surveillance period.

**Figure 3:**
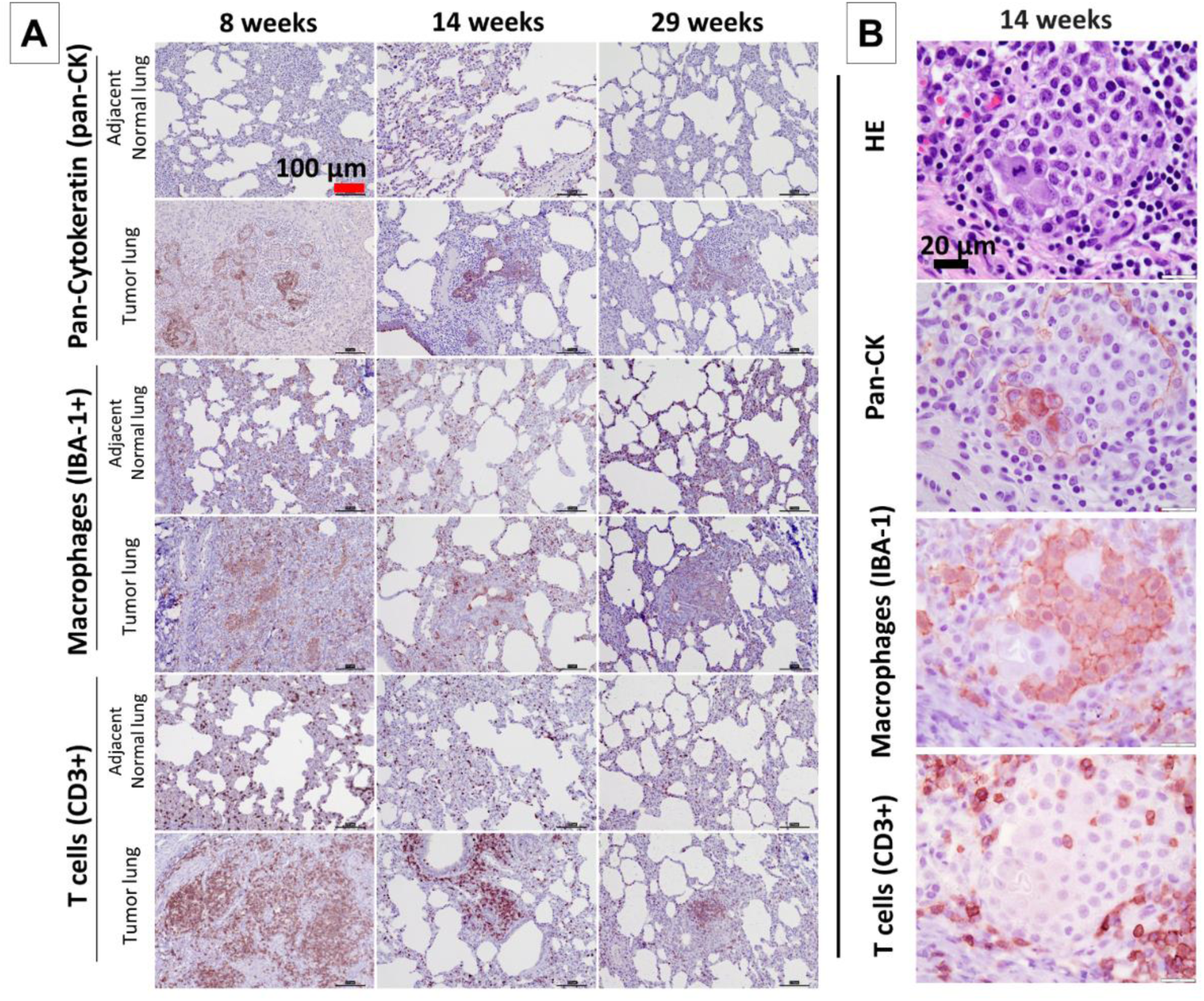
**A:** Immune cell infiltration in the Oncopig LC tumor microenvironment at three different autopsy dates at 8, 14 and 29 weeks. Immunohistochemistry staining for IBA-1 (macrophages) and CD3 (T cells) identified immune cells in LCs and adjacent normal lungs, while the extent of infiltration subsequently decreased. Pan-CK staining is also shown. (Scale bars, 100 μm.). **B**: **T cell and macrophage infiltration around invasive cancer cells in the tumor microenvironment of an Oncopig LC at higher magnification.** Histopathology (H&E) staining and immunostaining for identification of Pan-CK+ epithelial cells, in addition to IBA-1+ macrophages and CD3+ T cells in the tumor microenvironment are shown. (Scale bars, 20 μm).

### LC cell expression suggests cell proliferation and epithelial-mesenchymal transition (EMT) as biomarkers of malignancy and metastatic potential

Immunohistochemistry for proliferation and EMT-associated markers was also performed in Oncopig-derived LC tissues and matched soft tissue metastasis (**Figure 4**), showing decreased expression of E-cadherin in the soft tissue metastasis compared to the primary LC. Additionally, similar expression pattern of proliferation marker Ki-67 between primary LC and the soft tissue metastasis was noted, whereas higher expression of EMT-marker vimentin was observed in the soft tissue metastasis. These findings demonstrate cancer cell proliferation and metastasis-associated EMT marker expression patterns in Oncopig-derived LC tissues.

**Figure 4.**
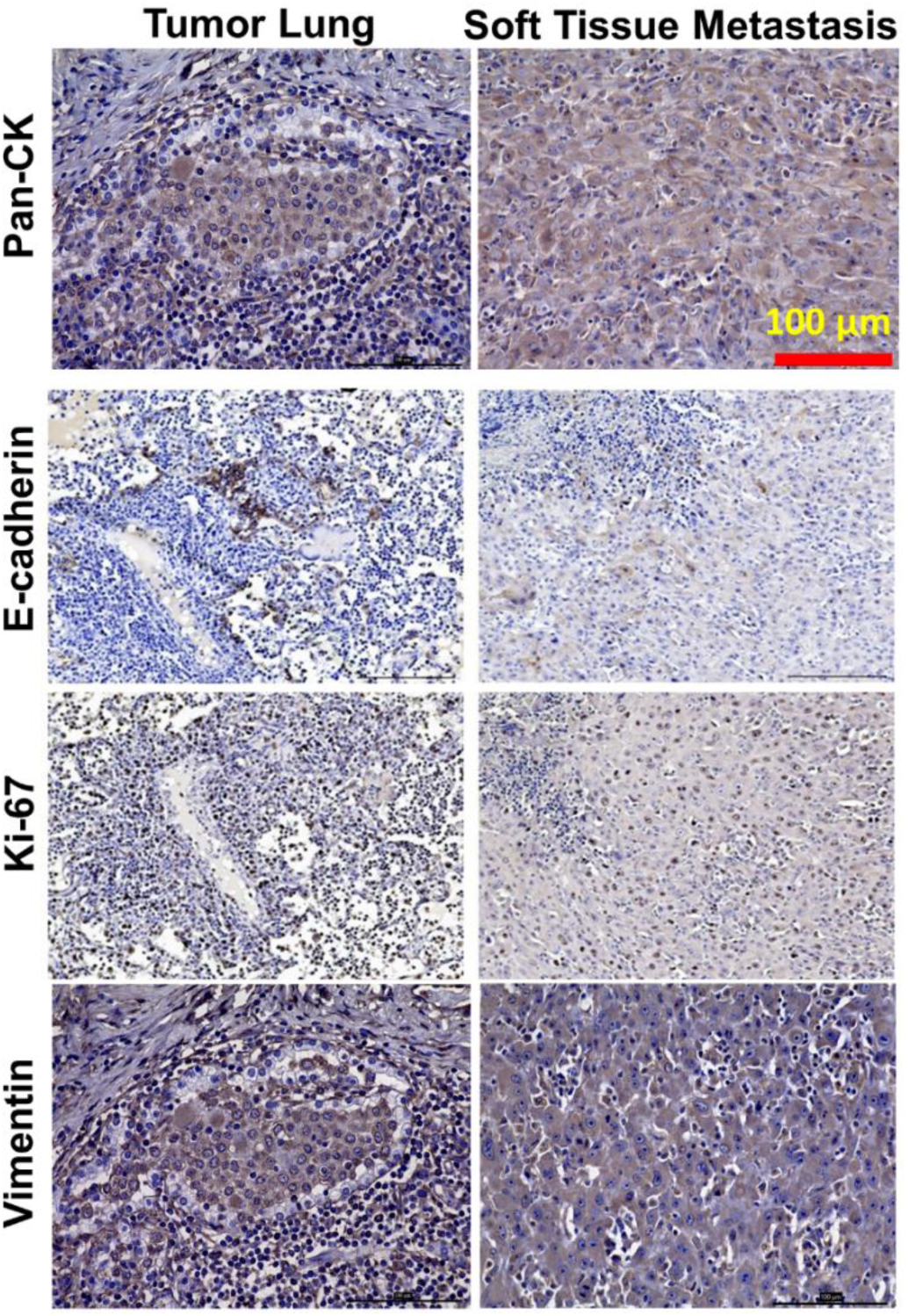
Oncopig-derived LC and soft tissue metastasis (matched) showing decreased expression of E-cadherin (2^nd^ row) in the metastasis. Nuclear expression of proliferation marker Ki-67 (3^rd^ row) and epithelial-to-mesenchymal transition (EMT) marker vimentin (4th row) was observed in both primary LC and the metastasis. Pan-CK expression is also shown (upper row). (Scale bars, 100 μm).

### Gene expression of Oncopig LC has significant overlap with human LC

Gene expression analysis of Oncopig-derived LC tumor tissues and normal Oncopig lung tissue (n=2, respectively) was performed by whole transcriptome sequencing to identify the molecular expression landscape, and also allow comparison with human LC patients’ data **(Figure 5**). Principal component analysis revealed distinct changes in the transcriptome between Oncopig tumor samples versus normal pig lung tissues (**Figure 5A**). Differential gene expression analysis showed overexpression of classic epithelial genes (e.g., SFTPC and STEAP1), genes associated with cancer (e.g., CLDN6), and regulators of EMT (e.g., vimentin, fibronectin 1, and N-cadherin) in Oncopig LC versus normal pig lung tissues (**Figure 5B**). These gene expressions demonstrate cancer-specific transcriptome changes in Oncopig LC. Further, gene ontology enrichment analysis of the top 50 differentially expressed genes between LC and normal tissues showed cell adhesion and migration pathway enrichment in tumor tissues over normal lung tissues (**Figure 5C**).

**Figure 5.**
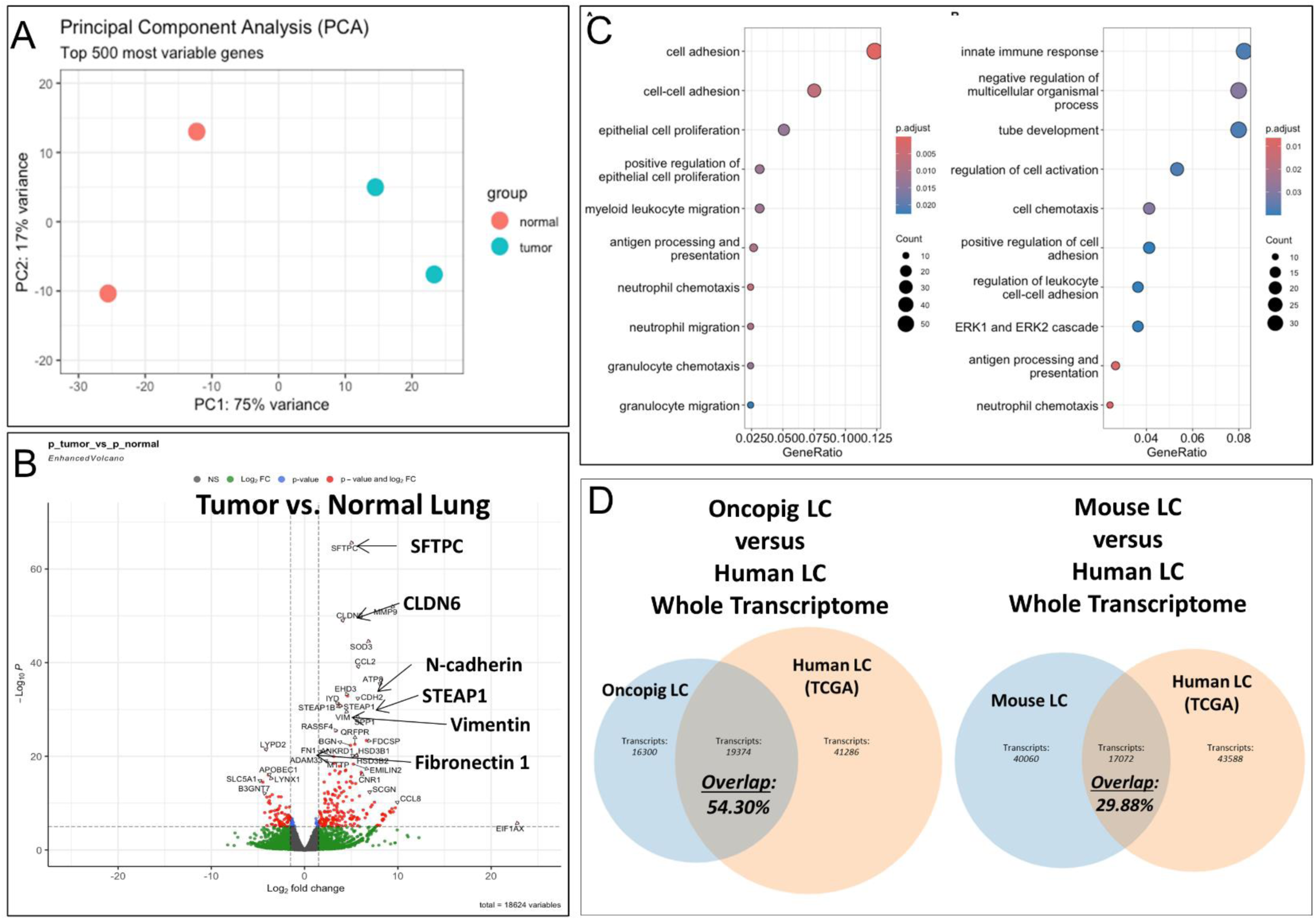
Whole transcriptome analysis of Oncopig lung tumors and healthy pig lung in comparison with human LC TCGA data. **A:** Principal component analysis of Oncopig-derived LC and normal Oncopig lung showing similarities and variabilities between normal and tumor tissues (n=2, respectively). **B:** Differential gene expression analysis between LC tumor versus normal pig lung tissues. **C**: Gene ontology and pathway enrichment analysis of top 50 differentially expressed genes of Oncopig LCs. **D:** Orthologous gene expression comparison with human LC patients was performed using The Cancer Genome Atlas (TCGA) data [total N=40 non-small cell lung cancer (NSCLC) patients’ transcriptomic data [N=10 per stage I-IV, respectively)], demonstrating 54.3% of the Oncopig LC transcripts overlapping with human LC transcripts, in contrast to less (29.88%) overlap with mouse KRAS-mutant LC.

For comparative analysis of Oncopig LC with human LC gene expression data, orthologous gene expression analysis was performed against The Cancer Genome Atlas (TCGA) [total N=40 non-small cell lung cancer (NSCLC) patients’ data (N=10 per stage I-IV, respectively)]. This transcriptome comparison of differentially expressed genes of pig and human LC revealed 54.3% overlap of Oncopig LC with human LC transcriptome (**Figure 5D; left panel**). A similar orthologous analysis revealed substantially less (29.88%) overlap between KRAS-mutant mouse LC and human LC (**Figure 5D; right panel**)

## DISCUSSION

Scientific findings in pigs are highly translatable to human patients ^2, 7, 26^ due to shared similarities such as in anatomy, physiology, metabolism ^3, 4, 6, 8, 27, 28^. Pigs also allow to use of human-grade instrumentations, which is of tremendous advantage for studying human diagnostic and therapeutic modalities including medical devices (e.g., ablation technologies), imaging/theranostics, radiation, and surgical techniques ^26^. To develop a robust preclinical LC model, we expanded the technical feasibility of a transgenic pig model for LC in a larger pig cohort with a longer surveillance period.

Transbronchial injection led to a successful tumor induction rate of invasive cancer and CIS in >90%, substantially higher than endobronchial injection in our cohort and endovascular/percutaneous injection in a recent pilot study ^21^. Of note, no procedure-associated complications were encountered with transbronchial needle injections that can be performed with minor operator training in a minimally invasive fashion using a flexible bronchoscope without the need for radiographic image guidance with radiation exposure. Also, transbronchial injections allowed a AdCre depot injection with presumable higher local concentration into a reproducible lung target area, rather than a more diffuse endobronchial or endovascular injection that may not accomplish sufficient AdCre concentrations for tumor induction. A percutaneous lung intervention, such as a percutaneous needle injection, has a significant risk of a pneumothorax due to violation of the visceral pleura which can also be avoided with a more central transbronchial needle injection of AdCre that does not violate the visceral pleura. Histopathologically, the LCs showed significant tumor microenvironment immune cell infiltration, opening avenues to study resistance mechanisms toward immunotherapy in LC. Oncopig LC cells also expressed cancer-associated biomarkers, with gene expression matching human NSCLC at a high rate of >54%.

The vast majority of the cancer treatment effects investigated in a variety of mouse models ^29–37^ have subsequently not been reproducible in human clinical trials ^2^. More recently, patient tissue-derived tumor organoids have evolved as promising drug testing platforms for personalized medicine ^38–40^, but all human-derived models are limited by lack of repeated tissue availability, loss of local tumor microenvironment, and inability to represent the systemic immune responses ^34^. So far, pig cancer models have also had challenges beyond more complex maintenance and higher costs. For instance, human cancer cell lines were xenografted into pigs ^41^ with a need for immunodeficient animals unsuitable for immunotherapy study ^42^. Other pigs were genetically engineered to carry a single, very specific, and often rare mutation without broader clinical applicability ^43^. In addition to being immunocompetent, the Oncopig ^10^ is transformative as it carries a most common mutant tumor suppressor (TP53^R167H^) and a commonly found (20-50%, depending on cancer type ^15^) oncogenic mutation (KRAS^G12D^) orthologous to human cancer mutations ^11, 12^ that can be induced orthotopically in the lung.

In a pilot study of four Oncopigs, LC has been induced differently by AdCre delivery with endovascular and percutaneous injection ^21^. The surveillance was 21 days, and neoplastic lung nodules developed in 10% of endovascular and in 33% of percutaneous injections ^21^. In our study, we expanded the cohort to 12 Oncopigs with longer surveillance and ultimately accomplished higher (>90%) LC induction via transbronchial needle injections. We confirmed epithelial cancer cells that expressed EMT and other cancer biomarkers, whereas the tumor microenvironment contained abundant macrophages and T cells ^21^. Of note, all carcinoma in situ (CIS) were observed following transbronchial injections with a higher AdCre dosage in group 3 that was followed comparably brief for 14 weeks, so that we may have likely seen a progression to invasive cancer if we had followed them longer. In one Oncopig, a soft tissue metastasis was identified incidentally on autopsy with histopathological confirmation. As described in the previous pilot study ^21^, we also found desmoplastic stroma and undifferentiated rather non-small cell cancers without typical characteristics of adenoid nor squamous cell subtypes of NSCLC and no features of small cell carcinoma (SCLC).

The inflammatory reaction consisting of macrophages and T cells in the LC tumor microenvironment could be of benefit to future studies of immunotherapy responses. Of note, the Oncopig was used for pancreatic cancer induction with a success rate of 71%, but the investigators also reported severe inflammation/pancreatitis ^22^. Oncopigs with pancreatic injections of adjuvant IL-8 without AdCre and wild-type pigs with AdCre injection did not show notable inflammation ^22^, so that the tumor-associated inflammation may be secondary to an immune response to an acute load of tumor-associated neoantigens ^21, 22^. In our study, we also did not observe any inflammatory reactions at the control injection sites with adjuvants without AdCre. An immune response from AdCre injection within the Oncopig skeletal muscle has been characterized as an intratumoral infiltration of cytotoxic T cells with enrichment of regulatory (FOXP3+) T cells with increased expression of immune checkpoint inhibitor targets indoleamine 2,3-dioxygenase 1 (IDO1), cytotoxic T-lymphocyte-associated protein 4 (CTLA4), and programmed death-ligand 1 (PDL1) ^44^. These findings were interpreted as a beneficial antitumor immune response and support the hypothesis that the Oncopig may serve as a valuable model to study tumor-directed cytotoxicity ^44^. Of note, in contrast to rodents the porcine immune system is much more similar to that of humans, again suggesting higher translatability ^45^.

Upon further molecular characterization for cancer hallmark biomarkers, with comparable results observed in the Oncopig pancreatic cancer model ^22^, we found downregulation of cell-cell adherens junction transmembrane protein E-cadherin and expression of EMT marker vimentin as indicators of metastatic capacity. Increased cell proliferation was also observed. In comparison to untransfected Oncopig lung tissue, there was significantly higher differential gene expression in Oncopig LC with regard to cell adhesion, proliferation, and EMT-associated genes. Importantly, transcriptome comparison with human TCGA NSCLC patient data showed a striking overlap of 54.30% matching with human LC transcriptome. We also performed a similar comparison ^46^ of LCs generated in KRAS-mutant mice^47^ that showed substantially less overlap (29.88%) of orthologous genes between mouse and human LC. Our findings in LC are consistent with previous cross-species transcriptome analyses between mice and humans that revealed a low proportion of ortholog transcripts of <20% ^48^, further supporting that findings in pigs are much more translatable to humans than findings in mice.

There are several limitations to consider in our pig study that was associated with higher costs and logistical efforts in comparison to rodents. Foremost, we did not achieve 100% induction success rates, although we accomplished higher cancer induction efficiency than in any other Oncopig cancer study with transbronchial injections leading to >90% of at least CIS. Due to the low sample size of twelve Oncopigs, we also allowed some interpretation bias as we administered different dosages of AdCre and modifications in adjuvants compositions to enhance chance of LC induction. There is limited availability of anti-porcine antibodies restricting extensive molecular and pathological characterization for subtyping of the Oncopig LCs. A specific consideration for LC is that the current Oncopig line carries the quite prevalent *KRAS^G12D^* mutation, yet it is more common in gastrointestinal cancers ^49^. The glycine-to-cysteine mutation in position 12 (*KRAS^G12C^*) is the most common *KRAS* mutation in LC (>15%) ^16^ with available FDA-approved drugs (e.g., sotorasib) ^50^. Clearly, an Oncopig line carrying KRAS^G12C^ would be most appropriate to study KRAS-associated drug resistance in LC.

## CONCLUSIONS

A transgenic, immunocompetent, and orthotopic LC Oncopig model appears highly innovative with expected clinical impact for human LC care. Preclinical testing in an Oncopig LC model (including bronchoscopic interventions, ablations, medical devices, radiation approaches, theranostics, targeted and immunotherapies) ^51^ could be impactful with a variety of modalities that are challenging or impossible to test in smaller animals. It is anticipated that a highly translatable pig LC model can lead to novel findings for precision oncology to improve LC patients’ outcomes.

## DECLARATIONS

## Acknowledgments

We are exceedingly grateful to the animal care staff at the Veterinary School, NextGen Precision Health Institute, and the National Swine Resource and Research Center (NSRRC) at the University of Missouri. We are also very grateful for the assistance of the staff from the National Swine Testing Center (U42OD035738) regarding helping with the animal experiments.

## Statement of ethics approval

The study was approved by the Institutional Animal Care and Use Committee at the University of Missouri (protocol # 18120). Animal housing facilities are accredited by AAALAC and in compliance with and regulated by USDA (43R0048) and Office of Laboratory Animal Welfare (D16-00249).

## Author contributions

J.K., K.N.S., N.S.N., Y.M., T.H., B.N., B.P.T., J.T.K., S.R.: conceptualization, conduction of the experiments, methodology data analysis, original manuscript draft writing; J.C.: histopathology, immunohistochemistry, pathology interpretation; K.N.S., Y.M., R.M., J.T.K: bioinformatic analysis of transcriptome data; J.R.K.: conductance and interpretation of radiographic imaging; M.G., R.M.: statistical analysis; R.S.P., S.S., K.W., J.N.B., E.T.: conceptualization, methodology, interpretation of data, review & editing. All authors reviewed and approved the submission of the manuscript.

## Availability of data and materials

All data generated or analyzed during this study, if not included in this article and its supplementary information files, are available from the corresponding authors on request.

## ABBREVIATIONS

AAALAC: Association for Assessment and Accreditation of Laboratory Animal Care International

AdCre: Adenovirus Cre

ANOVA: Analysis of Variance

BMP: Basic Metabolic Panel

CBC: Complete Blood Counts

CK: Cytokeratin

CT: Computed Tomography

CRE: Cyclic Recombinase

DMSO: Dimethylsulfoxide

DNA: Deoxyribonucleic acid

FDA: United States Food and Drug Administration

IHC: Immunohistochemistry

KRAS: Kirsten Rat Sarcoma Virus gene

LC: Lung Cancer

NSRRC: National Swine Resource and Research Center

PFF: Porcine Fetal Fibroblasts

RNP: Ribonucleoprotein

RNA: Ribonucleic Acid

SD: Standard deviation

SEM: Standard error of the mean

TCGA: The Cancer Genome Atlas

TP53: Tumor Protein P53

