## Supplementary Methods for "Characterization of A Bronchoscopically Induced Transgenic Lung Cancer Pig Model for Human Translatability"

### SUPPLEMENTARY SECTION

#### Whole transcriptome sequencing and bioinformatic analysis

For whole transcriptome analysis of Oncopig-derived LC tumor (n=2) and normal lung tissues (n=2), RNA extraction was done with cellular lysis using MagMax mirVana Total RNA Isolation kit (A27828; Thermo Fisher), following the manufacturer's instructions. The RNA samples were subjected to whole transcriptome analysis (RNA sequencing) at ArrayStar Inc. (Rockville, MD). Assessment of RNA quality was accomplished using NanoDrop ND-1000 instrument. ~1-2 µg total RNA was used to prepare the sequencing library. Briefly, total RNA was enriched by oligo (dT) magnetic beads (rRNA removed) followed by RNA-seq library preparation using KAPA Stranded RNA-Seq Library Prep Kit (Illumina). The completed libraries were qualified with Agilent 2100 Bioanalyzer and quantified by absolute quantification qPCR method. To sequence the libraries on the Illumina NovaSeq 6000 instrument, the barcoded libraries were mixed, denatured to single-stranded DNA in NaOH, captured on Illumina flow cell, amplified in situ, and subsequently sequenced for 150 cycles for both ends on Illumina NovaSeq 6000 instrument.

*Pre-processing and normalization*: The fastq reads obtained were preprocessed to remove low-quality reads and adapters using fastp (v0.02.0). Quality control was performed using FastQC (v0.11.9). The clean reads were aligned to the Pig reference genome (Sscrofa11.1) using STAR (v2.7.10) with default parameters. Gene expression levels were quantified, and the resulting count data were normalized for differences in library size by estimating size factors.

*Principal Component Analysis (PCA)*: PCA was conducted to examine overall similarities and variations between experimental groups. The PCA plot was generated using the plot PCA function by plotting the first two principal components on the x and y axes. The top 500 most variable genes were used for the PCA plot.

*Differential Gene Expression Analysis*: DESeq2 R package (v1.42.0) was used to find differentially expressed genes (DEGs). To estimate mean expression levels between normal and tumor samples, DESeq2 with a negative binomial generalized linear model and Wald test was used to evaluate the significance of log2 fold changes. Multiple testing correction for controlling the false discovery rate (FDR) was applied using the Benjamini-Hochberg procedure. Genes were identified as differentially expressed based on adjusted p-values < 0.05 and log2 fold changes ±1.5. A volcano plot was generated to visualize the relationship between fold change and the statistical significance of DEGs. The enhanced Volcano function from the R package (v1.20.0) was used to generate the plot, highlighting significant DEGs with an adjusted p-value of less than 0.05.

*Gene Ontology (GO) Enrichment Analysis*: GO enrichment analysis was performed to identify significantly overrepresented genes to understand the enriched biological processes. The analysis involved calculating the expected and observed frequencies of GO terms and applying statistical tests (Fisher's exact test) to determine significant overrepresentation. Results were visualized as dot plots and cluster plots using the clusterProfiler R package.

Human and OncoPig Cross-Species Genome Comparison: A comprehensive comparative transcriptomic analysis was conducted to identify and compare gene expression patterns between OncoPig and human tumors. Human NSCLC transcriptome RNA-seq count data and clinical pathological information, including stage, were acquired from The Cancer Genome Atlas (TCGA) database. The data were downloaded in raw count format. Initial preprocessing involved assessing data quality, filtering out low-quality reads, and ensuring data integrity. The raw counts were further processed using the DESeq2 R package (v1.42.0) to normalize the data and mitigate variations in sequencing depth and RNA composition among samples. Orthologous gene mapping between OncoPig and human genomes was performed using orthogene interspecies gene mapping ensuring accurate identification of corresponding genes across species. Following this, gene expression data from both species were integrated and normalized to account for differences in sequencing depth and RNA composition using the DESeq2 framework. The normalized datasets enabled a robust comparative analysis to pinpoint conserved and divergent gene expression features relevant to lung cancer. Similar analysis was performed using publicly available KRAS-mutant mouse lung cancer datasets to identify the overlap of differential expressed genes between mouse and human lung cancer. Briefly, KRAS-mutant mouse lung tumor (n=3) and normal lung (n=3) RNA-seq FastQ files were downloaded from NCBI gene expression omnibus using BioProjects PRJNA1033182 and PRJNA1090424, respectively. FastQ reads were preprocessed and the resulting count data was used as described above to identify orthologous gene expression overlap between mouse and human lung cancer differentially expressed genes.
